# NAD^+^ metabolism is a key modulator of bacterial respiratory epithelial infections

**DOI:** 10.1101/2023.04.13.536709

**Authors:** Björn Klabunde, André Wesener, Wilhelm Bertrams, Isabell Beinborn, Nicole Paczia, Kristin Surmann, Sascha Blankenburg, Jochen Wilhelm, Javier Serrania, Kèvin Knoops, Eslam M. Elsayed, Katrin Laakmann, Anna Lena Jung, Andreas Kirschbaum, Mobarak Abu Mraheil, Anke Becker, Uwe Völker, Evelyn Vollmeister, Birke J. Benedikter, Bernd Schmeck

## Abstract

Lower respiratory tract infections caused by *Streptococcus*O*pneumoniae (Spn)* are a leading cause of death globally. Here we investigate the bronchial epithelial response to *Spn* infection on a transcriptomic, proteomic and metabolic level. We found the NAD^+^ salvage pathway to be dysregulated upon infection in a cell line model, primary human lung tissue and *in vivo* in rodents, leading to a reduced production of NAD^+^. Knockdown of NAD^+^ salvage enzymes (NAMPT, NMNAT1) increased bacterial replication. NAD^+^ treatment of *Spn* inhibited its growth while growth of other respiratory pathogens improved. Boosting NAD^+^ production increased NAD^+^ levels in immortalized and primary cells and decreased bacterial replication upon infection. NAD^+^ treatment of *Spn* dysregulated the bacterial metabolism and reduced intrabacterial ATP. Enhancing the bacterial ATP metabolism abolished the antibacterial effect of NAD^+^. Thus, we identified the NAD^+^ salvage pathway as an antibacterial cascade in *Spn* infections, predicting a novel antibacterial mechanism of NAD^+^.

## 2. Introduction

Lower respiratory infections are the fourth leading cause of death globally, with *Streptococcus*□*pneumoniae* (*Spn*) being the most common respiratory bacterial pathogen (GBD 2019 Antimicrobial Resistance Collaborators, 2022). Thus, an in-depth understanding of infection processes on the molecular level is vital to enabling the development of new therapeutic options to face rising antibiotic resistance. Accordingly, host–pathogen interactions during pneumococcal infections are a field of intensive study. However, while most studies have focused on host responses towards *Spn* in dedicated immune cells (Ritchie and Evans, 2019), the respiratory epithelium constitutes the first line of defense against lower respiratory tract infections. It forms a mechanical barrier, produces a thick, glycan-composed mucus to entrap bacteria and secretes various cytokines and chemokines to recruit immune cells to sites of infection. It also promotes direct bacterial killing by secreting antibacterial peptides (Kuek and Lee, 2020). Analyzing the immediate transcriptomic response of infected alveolar epithelial cells up to 4 h post *Spn* infection, Aprianto *et al*. have demonstrated an extensive dysregulation of gene expression, especially repression of genes involved in the immune response (Aprianto *et al*., 2016). Yet, the effects of prolonged *Spn* infection on epithelial cell gene expression and metabolism have not yet been investigated.

Here, we employed a multi-omics approach to gain insights into the epithelial response against *Spn* infection. This approach revealed a dysregulation of key enzymes and metabolites of the Nicotinamide adenine dinucleotide (NAD) metabolism. By transitioning between its oxidized (NAD^+^) and reduced (NADH) forms, NAD acts as key electron transporter in cellular energy pathways. In addition, it is an important cofactor for a wide array of enzymes that convert NAD^+^ to the energy depleted metabolite nicotinamide (NAM) (e.g., PARP, sirtuins (Imai et al., 2000; Katsyuba et al., 2020; Wilk et al., 2020)). *De novo* biosynthesis of NAD^+^ starts from diet-derived tryptophan or nicotinic acid (Yang and Sauve, 2016). However, the majority of cellular NAD^+^ is recovered from NAM via the NAD^+^ salvage pathway (Strømland et al. 2019). In a first salvage step, nicotinamide phosphoribosyl transferase (NAMPT) catalyzes the production of nicotinamide mononucleotide (NMN) from NAM (Revollo et al., 2004). Second, nicotinamide-mononucleotide adenylyl transferases (NMNAT) use NMN as substrate to restore NAD^+^ (Sestini et al., 2000; Strømland et al., 2019).

To explore the functional relevance of the altered NAD^+^ metabolism described here, we manipulated different enzymes or metabolites and explored their effects on *Spn* infection. Thereby, we identified the modulation of the NAD^+^ metabolism as a previously unknown element in antibacterial defense against *Spn*. Direct treatment of *Spn* with NAD^+^ inhibited bacterial growth in a concentration-dependent manner, while other bacterial pathogens remained unaffected. Our study deepens the understanding of the interaction between *Spn* and the host metabolism and identifies the NAD^+^ metabolism as a potential therapeutic target against lower respiratory tract infection.

## 3. Results

### 3.1. Pneumococcal infection dysregulates the epithelial NAD^+^ metabolism

To investigate host–pathogen interactions during *Spn* infection of epithelial cells, we established an *in vitro* infection system using the bronchial epithelial cell line BEAS-2B (Figure S1). To assess the proteomic response of epithelial cells to *Spn* by SILAC proteomics, BEAS-2B cells were infected with MOI 0.5 for 16 h or left uninfected. (Figure 1A). Strikingly, the 5 most significantly upregulated proteins included two enzymes involved in the NAD^+^ metabolism, Nicotinamide phosphoribosyl transferase (NAMPT; fold change (FC) = 2.6, p=0.000002) and Nicotinamide N-Methyl transferase (NNMT; FC=1.7, p=0.000148). Significant regulation of the NAD^+^ metabolism was further confirmed by mRNA microarray data (Figure 1B). NAMPT and NNMT mRNA were upregulated compared to untreated controls after both *Spn* infection and stimulation with the bacterial cell wall component lipoteichoic acid (LTA) for 9 or 16 h. In addition, mRNA of nicotinamide mononucleotide adenylyl transferase 1 (NMNAT1) was significantly downregulated exclusively during later-stage infection (16 h). Differential mRNA regulations were confirmed by qPCR (NAMPT, NNMT, NMNAT1) and Western Blot (NAMPT; Figure S3). Metabolite measurements by LC-MS/MS revealed that NAD^+^, as well as multiple NAD^+^ precursors, were decreased after 16 h of *Spn* infection (Figure 1C). In summary, we found a clear dysregulation of NAD^+^ biosynthesis via the Preiss-Handler and NAD^+^ salvage pathways on a multi-omics scale (Figure 1D, Figure S2).

**Figure 1.**
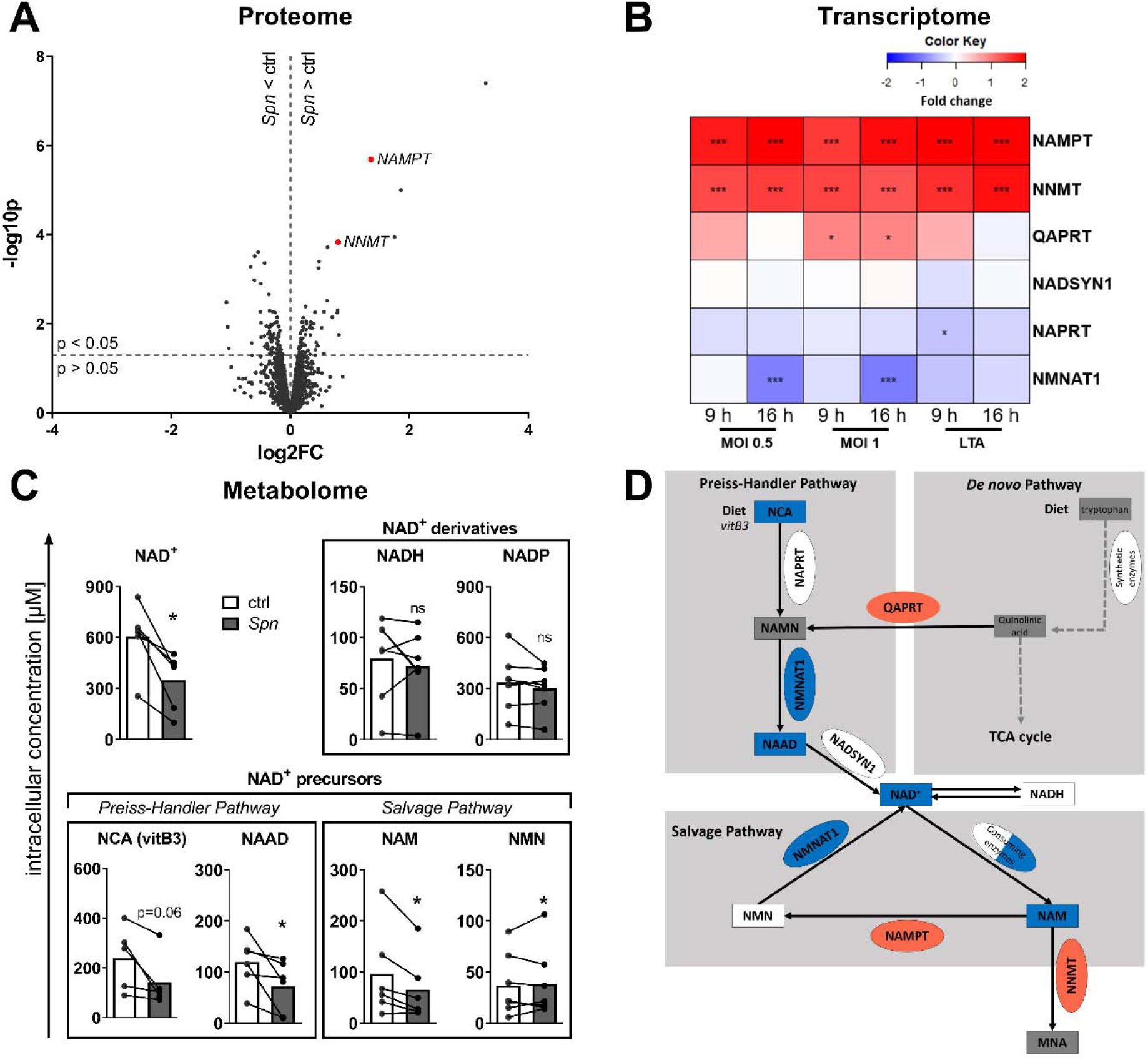
NAD^+^ biosynthesis is dysregulated during Spn infection. A) Volcano plot of protein expression upon Spn infection. BEAS-2B cells were infected with Spn MOI 0.5 for 16 h or left uninfected (ctrl). Afterwards, proteins were isolated and changes in protein expression were determined by SILAC proteomics. Regulated proteins associated with the NAD^+^ metabolism are marked in red. B) BEAS-2B were either left untreated, infected with Spn MOI 0.5 or 1, or treated with 1 µg/ml LTA for 9 or 16 h, respectively. RNA was isolated and gene expression was determined by mRNA microarray. NAD^+^ metabolism-associated genes are depicted as a heat map (value = fold change). C) BEAS-2B were infected with Spn MOI 1 for 16 h or left untreated. Metabolites were isolated and analyzed by LC-MS/MS. D) Summary of identified regulations of the NAD^+^ metabolism (red: upregulation; blue: downregulation; square: metabolite; oval: enzyme). Statistics: A) moderated t-test B) unadjusted p-values; C) paired t-test (* = p < 0.05;*** = p < 0.001; N = 3–6).

We then aimed to confirm the regulation of NAMPT and NMNAT1 upon pneumococcal infection in primary cell culture infection models. Primary human bronchial epithelial cells (hBECs) from healthy donors were differentiated into a pseudostratified lung epithelium at an air-liquid interface. Cells were apically infected with *Spn*, MOI 20 for 16 h and gene expression analyzed by qPCR (Figure 2A). In infected hBECs, we found expression of NAMPT to be upregulated and NMNAT1 to be downregulated compared to uninfected controls (Figure 2B, C). In a next step, human lung explants were injected with *Spn* for 12 h and gene expression determined (Figure 2D). We found an upregulation of NAMPT and downregulation of NMNAT1, confirming our previous results (Figure 2E, F). Re-analysis of publicly available gene expression datasets of *Spn* infected mouse lungs equally revealed NAMPT up-regulation and NMNAT1 downregulation *in vivo*. (Figure 2G, H). In summary, we found the NAD^+^ salvage gene expression to be dysregulated upon Spn infection in primary cell culture and *in vivo*.

**Figure 2.**
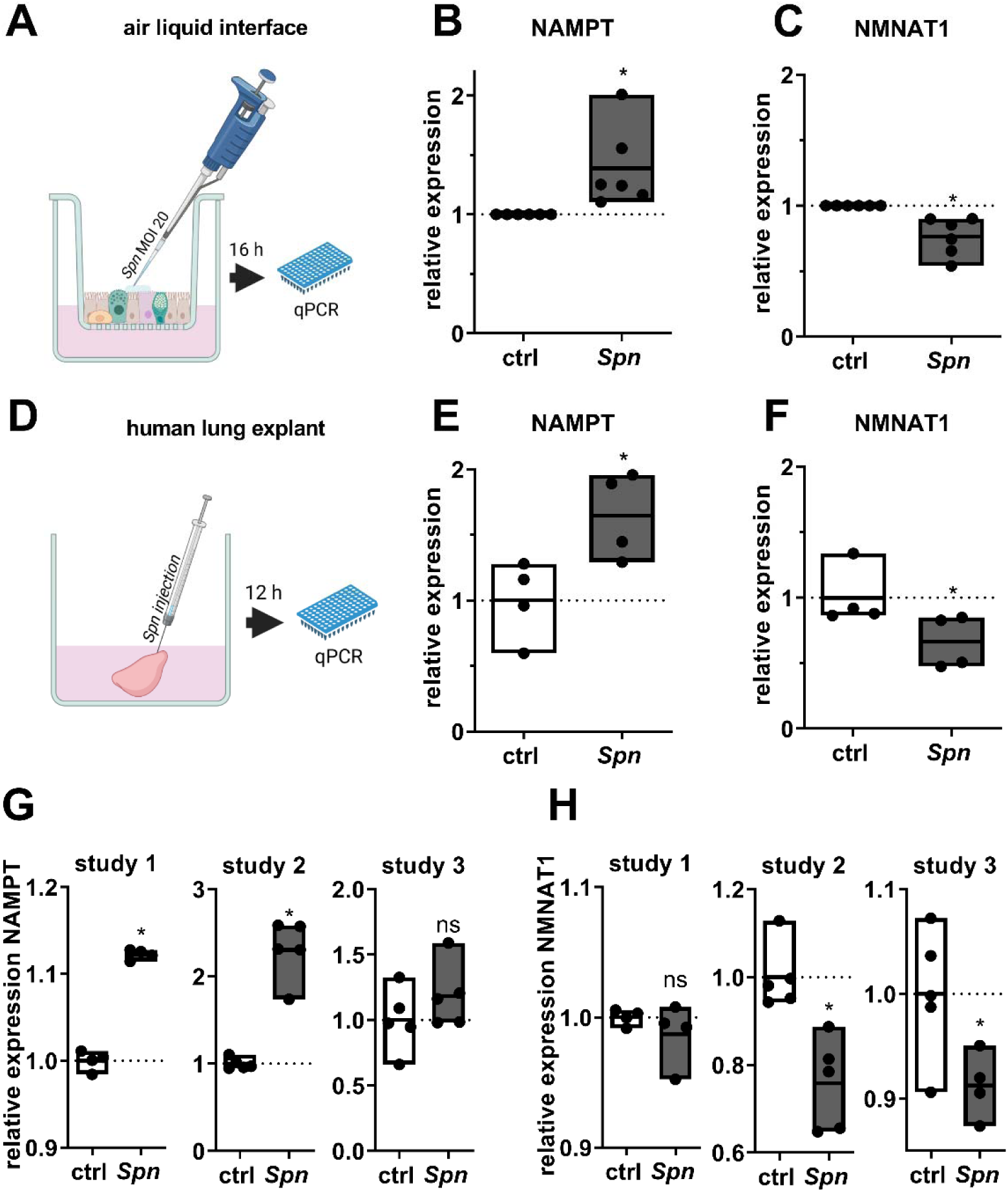
Regulation of NAD^+^ salvage associated genes ex and in vivo. A-C) Primary human bronchial epithelial cells from healthy donors were differentiated into pseudostratified respiratory epithelium as air–liquid interface cultures. After differentiation, cells were apically infected with Spn, MOI 20 for 16 h. The expression of NAMPT B) and NMNAT1 C) was determined by qPCR. (D-F) Human lung tissue explants were injected with Spn or PBS and incubated for 12 h D). Expression of NAMPT (E) and NMNAT1 (F) was determined by qPCR. G, H) 3 published microarray transcriptome datasets of murine lungs infected with different strains of Spn were analyzed for the expression of NAMPT (G) and NMNAT1 (H). Study 1: GSE83612, Mouse strain: C57BL/6, 24 h post-inoculation with 5 x 10^6^ CFU, serotype: 19; Study 2: GSE45644, mouse strain: BALB/C, serotype 2, 48 h post-inoculation with 5 x 10^4^ CFU,; Study 3: GSE61459, mouse strain: BALB/C, serotype: 2, 24 h post-inoculation with 5 x 10^6^ CFU. Statistics: paired t-test (B, C, E, F; t-test (G, H). (* = p < 0.05; n = 4-6 (B, C, E-H)); Results are normalized to untreated controls of the corresponding donor (C, D) or average expression values of control tissue/animals (E-H).

### 3.2. NAD^+^ production shows direct antibacterial effects

To assess the role of NAD^+^ biosynthesis during the infection process, we performed siRNA knockdowns of NAMPT and NMNAT1 48 h prior to infection with *Spn* (MOI 1, 16 h). Successful knockdowns were confirmed on transcript and protein level (Figure S4). After both knockdowns, intracellular NAD concentration was significantly decreased, whereas bacterial CFU after 16 h of infection was increased by approximately 40% compared to the scramble control (Figure 3A, B). Based on these findings, we further addressed whether the antibacterial activity of NAMPT and NMNAT1 is mediated by NAD^+^ production. Addition of NAD^+^ to the *Spn* infection of BEAS-2B cells (Figure 3C; MOI 1, 16 h) or to host cell-free cultures of *Spn* (Figure 3D-F) revealed a concentration-dependent reduction of bacterial replication, demonstrating a direct antibacterial effect of NAD^+^.

**Figure 3.**
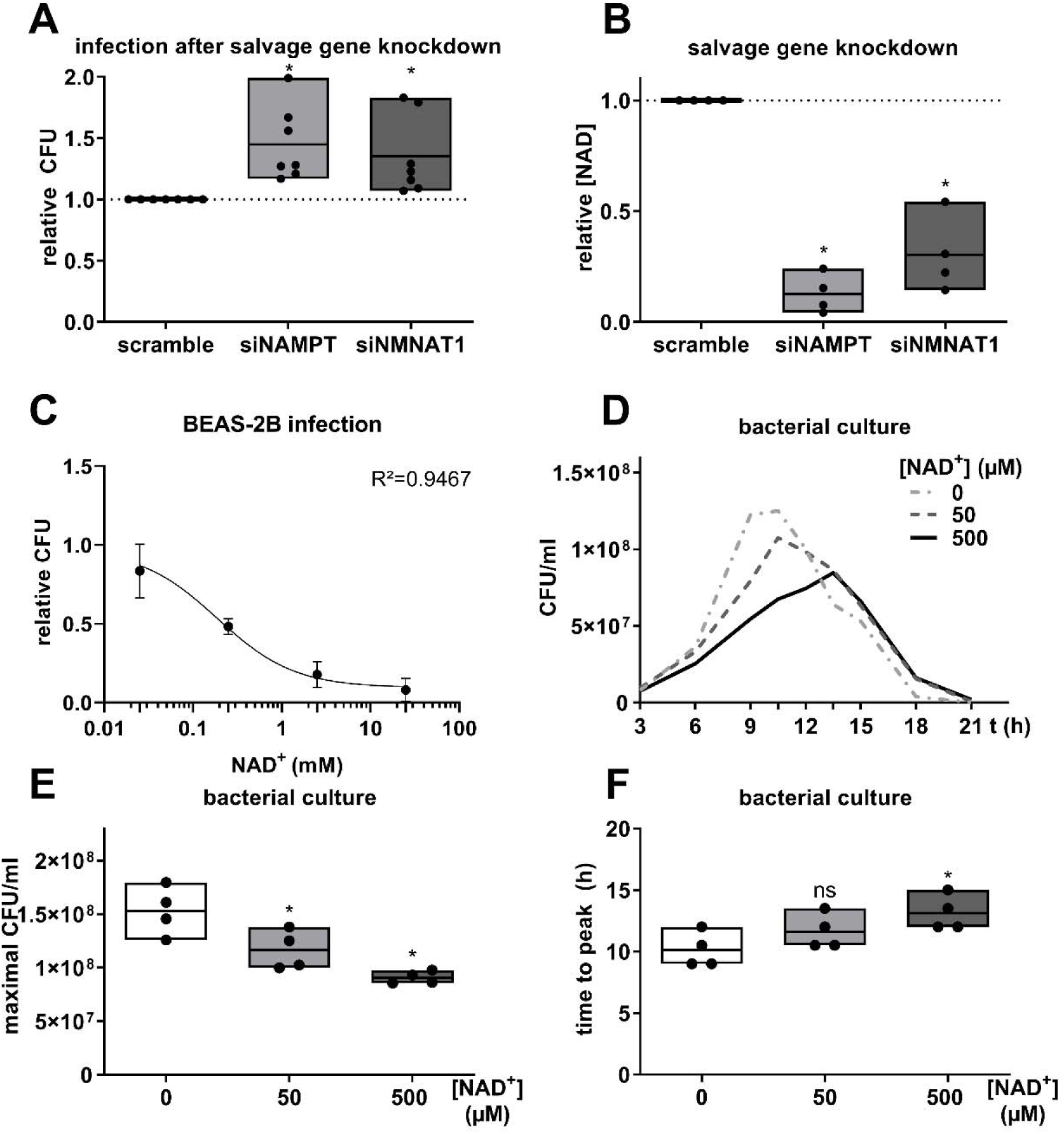
Effects of Nicotinamide metabolites on pneumococcal replication. A) BEAS-2B lung epithelial cells were transfected with indicated siRNA for 48 h prior to infection with Spn, MOI 1 for 16 h. Bacterial replication was determined by CFU assay. B) BEAS-2B cells were transfected as indicated for 48 h and the intracellular NAD content determined. C) BEAS-2B cells were infected with Spn, MOI 1 for 16 h. Indicated amounts of NAD^+^ were added at the start of the infection. Bacterial replication was determined by CFU assays 16 h post infection. D-F) Spn was cultivated in host cell-free medium with indicated amounts of NAD^+^. CFU were determined at indicated time points. D) Bacterial growth curves were measured over 21 h. The graph summarizes the mean of four biological independent replicates. E) Maximally reached CFU/ml of (D). F) Time (in h) until maximal CFU/ml was reached. Statistics: One-way ANOVA with Fisher’s LSD (A, B, E, F); Agonist vs Response-Fit (three parameters) (C); Significances were determined against scramble controls (A, B) or untreated controls (C, E, F) unless indicated otherwise. (ns = not significant; * = p < 0.05; N = 4-7).

To further assess the role of NAD^+^ during the infection process, we aimed to increase the host cell NAD^+^ production during infection. First, we boosted NAD^+^ production by treatment with the precursor NMN. NMN treatment of BEAS-2B increased the intracellular amount of NAD by a factor 2.5 compared to untreated controls (Figure 4A). In presence of host cells, NMN treatment reduced bacterial replication by factor 0.4 compared to untreated controls (Figure 4B). Strikingly, the effect was dependent on the presence of host cells, as NMN treatment of *Spn* cultivated in host cell-free medium did not affect bacterial replication (Figure 4B). Knockdown of NMNAT1 prior to infection and NMN treatment reduced NAD production (Figure 4C) and partially rescued bacterial replication (Figure 4D). Furthermore, chemical inhibition of NAMPT using FK-866 increased bacterial replication (Figure 4E), while NAMPT activation using SBI-797812 decreased *Spn* replication (Figure 4F). Next, we employed well-differentiated primary human bronchial epithelial cells at the air liquid interface model to confirm the effects of NMN addition on the course of pneumococcal infection (Figure 4G-I). During *Spn* infection, basolateral addition of NMN caused an increase in intra-and extracellular NAD (Figure 4G, H) and a reduction of bacterial replication (Figure 4I). In summary, knockdown and chemical modulation of the NAD^+^ biosynthesis enzymes NMNAT1 and NAMPT, as well as metabolite treatment of *Spn,* revealed an anti-pneumococcal effect of NAD^+^.

**Figure 4.**
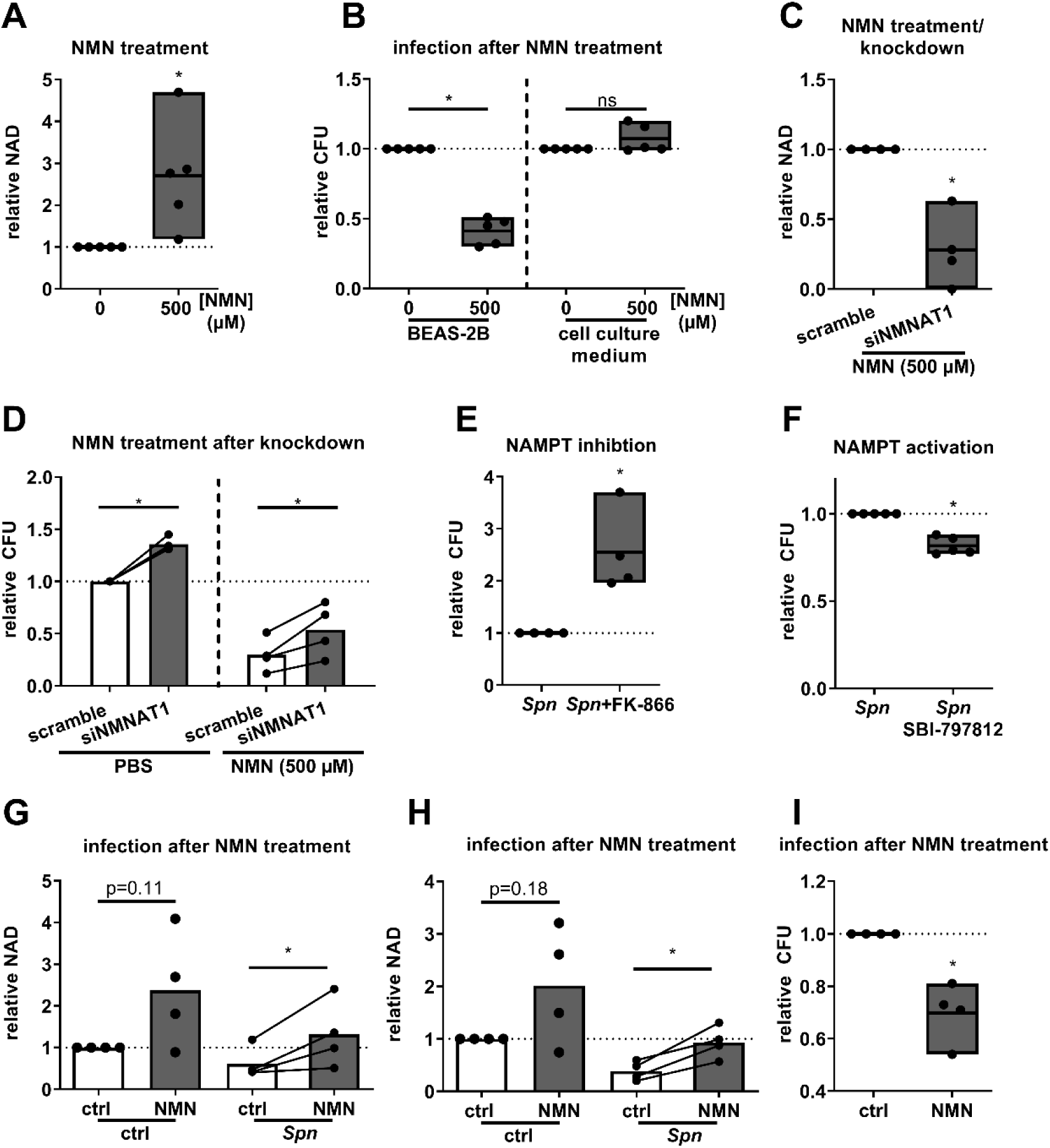
NMN-mediated growth inhibition is dependent on NAD^+^ production. A) BEAS-2B were treated with NMN for 9 h as indicated and intracellular NAD^+^ was measured. B) Spn were cultured in cell-free medium or used to infect BEAS-2B at MOI 1 with and without NMN treatment. CFU were determined after 9 h and are depicted relative. C) BEAS-2B were transfected with siNMNAT1 or a scramble control for 48 h. Afterwards, cells were treated with NMN for 9 h and the intracellular concentration of NAD was determined. D) Cells were transfected as described. 48 h post-transfection, cells were infected with Spn, MOI 1 or left uninfected and were treated with NMN (500 µM) or PBS. CFU/ml were determined 9 h post infection and normalized to scramble control. E) Cells were treated with the NAMPT inhibitor FK-866 (1 mM) 24 h prior to and during infection or left untreated. Bacterial CFU were determined 16 h post-infection. F) Cells were treated with the NAMPT activator SBI-797812 8 h prior to and during infection or left untreated. Cells were infected with Spn, MOI 1 for 16 h and CFU were determined. G-I) Primary pseudostratified human bronchial epithelial cells were cultivated at an air–liquid interface. Cells were infected apically with Spn, MOI 20 for 1 h. Afterwards, cells were washed, NMN (500 µM) was added basolateral and cells were incubated for 16 h. G, H) The amount of intracellular and extracellular NAD was determined 16 h post-infection in cell lysates (G) and apical wash fluid (H), respectively. I) Cells were washed 16 h post infection and CFU in the apical wash fluid were determined. Statistics: paired t-test; Significance was determined against uninfected controls unless indicated otherwise. (ns = not significant; * = p < 0.05; N = 4–5).

### 3.3. The antibacterial effect of NAD^+^ is specific to *Spn*

To determine whether interference with the host NAD^+^ metabolism is a general process affecting bacterial infections, we performed BEAS-2B infections with *Streptococcus agalacticae* (*S.aga*) and non-typeable *Haemophilus influenzae* (NTHi) and determined the resulting expression of NAMPT and NMNAT1 (Figure S5). Both bacteria induced a significant upregulation of NAMPT expression 16 h post-infection (Figure S5A). *S.aga* infection induced a significant downregulation of NMNAT1 by approximately a factor of 0.75, while NTHi infection, by tendency, caused an upregulation of NMNAT1 (Figure S6B). Next we investigated the effect of NAD^+^ treatment on *S.aga*, NTHi and an alternative, highly pathogenic strain of *Spn* (TIGR4). Treatment of *Spn* TIGR4 with NAD^+^ for 9 h caused a reduction in bacterial growth to approximately factor 0.4, confirming our previous results with *Spn* D39. In contrast, replication of *S.aga* and NTHi was increased by approximately 50% (Figure S5 D-F). In summary, these results reveal that, while the upregulation of NAMPT is a general pro-inflammatory process, downregulation of NMNAT1 expression and antibacterial effects of NAD^+^ appear specific to certain bacteria.

### 3.4. Transcriptional regulation of NMNAT1 is induced by *Spn* virulence factor pneumolysin

We next focused on the specific mechanisms by which *Spn* mediates NAD^+^ salvage gene regulation in epithelial cells. Upregulation of NAMPT was previously shown to be induced via JAK-STAT signaling under pro-inflammatory conditions (Dantoft et al., 2019). While we were could confirm JAK-dependency of NAMPT upregulation in response to *Spn* infection using the JAK inhibitor Ruxolitinib (Figure S6A), repression of NMNAT1 was not JAK-dependent (Figure S6B). Treatment of epithelial cells with heat-or UV-inactivated *Spn* did not influence the expression of NMNAT1, suggesting that the repression is actively mediated by *Spn* (Figure S6C, D). Therefore, we investigated the effects of two virulence factors which are actively produced and released during pneumococcal growth, the oxidant H_2_O_2_ and the pore forming toxin pneumolysin (*ply*) on NMNAT1 expression. Treatment with H_2_O_2_ did not influence NMNAT1 or NAMPT expression (Figure S6C, D). Infection with a *ply* deficient mutant of *Spn* D39 induced NAMPT gene expression to a similar extent as wild-type *Spn* (Figure 5B). In contrast, a *ply* deletion resulted by tendency in a reduced expression of pro-inflammatory IL-8 (Figure 5A) and in a significantly attenuated NMNAT1 repression compared to the wild type (Figure 5C). Treatment of BEAS-2B with purified, lytically active pneumolysin at non-lytic concentrations caused a dose-dependent pro-inflammatory response as indicated by the upregulation of IL-8 and a downregulation of both, NAMPT and NMNAT1 (Figure 5D-F). Interestingly, treatment with a lytically inactive, non pore-forming variant of pneumolysin did not cause a reduction of NAMPT and NMNAT1 expression (Figure 5 G-I). In summary, pneumolysin seems to mediate the reduction of expression NMNAT1 via pore formation.

**Figure 5.**
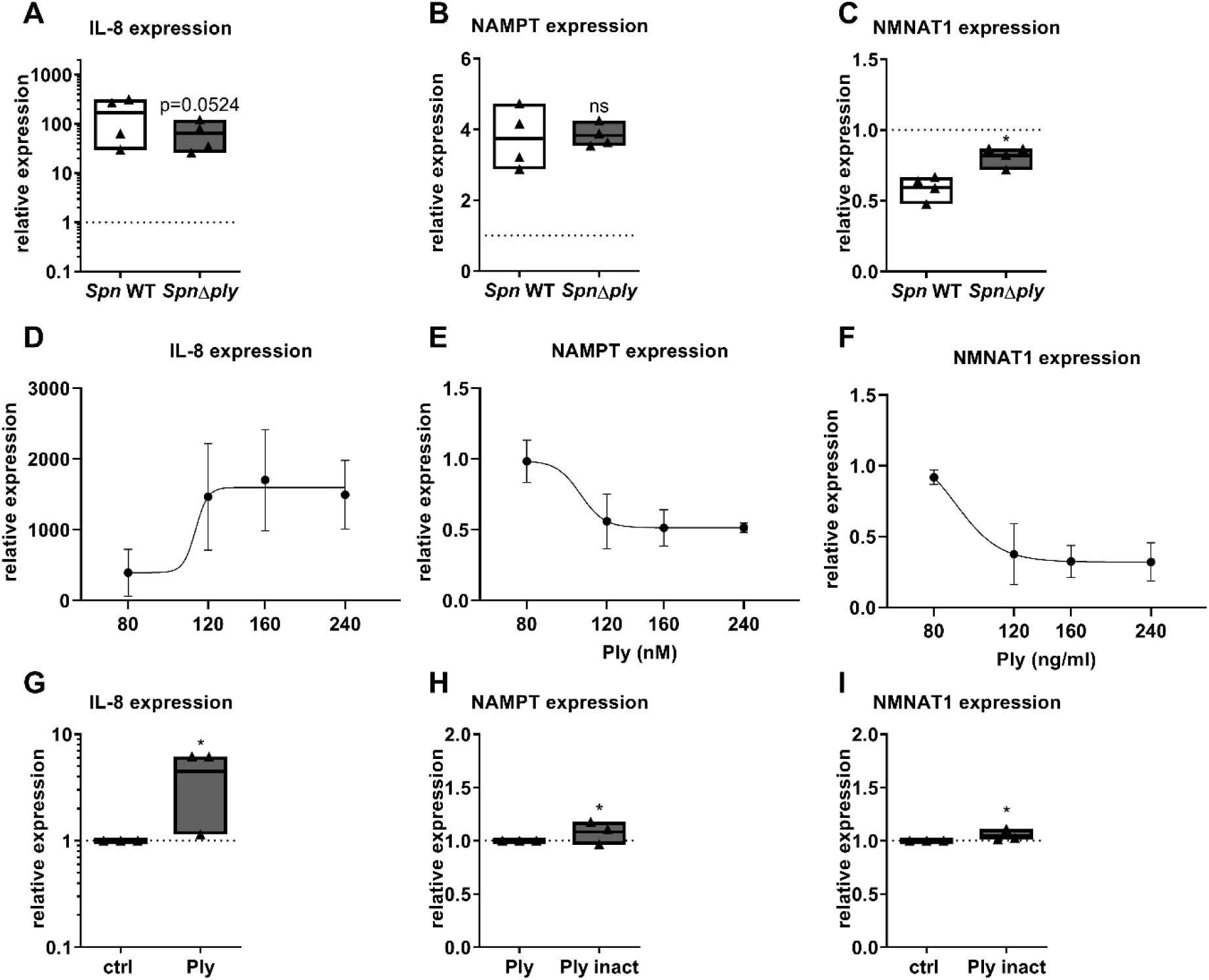
Transcriptional regulation of NMNAT1 is induced by pneumolysin (ply). A-C) BEAS-2B cells were infected with Spn, WT or Δply for 16 h. Gene expression was determined by qPCR. D-F) BEAS-2B cells were treated with pneumolysin in indicated concentrations for 16 h. Expression of IL-8, NAMPT and NMNAT1 was determined as indicated. Statistics: paired t-test (A-C, G-I); sigmoidal dose-response fit with variable slope (D-F); Significances were determined against uninfected or unstimulated controls unless indicated otherwise. (* = p < 0.05; ns = not significant; N=3-4).

### 3.5. Resistance against NAD^+^ is associated with a loss of capsule

We next investigated the antibacterial mechanisms of NAD^+^. First, we confirmed that extrinsic addition of NAD^+^ to *Spn* cultures results in increased total intrabacterial NAD (Figure 6A). Then, resistant clones were generated by cultivating *Spn* D39 in liquid medium with increasing concentrations (50 µM to 5 mM) of NAD^+^. After six passages, bacteria were plated on blood agar supplemented with 500 µM NAD^+^ and three clones were picked. All three clones were almost completely resistant against antibacterial effects of NAD^+^ (Figure 6B). Strikingly, genome sequencing revealed the capsule biosynthesis-associated gene *cps2E* to be mutated in all resistant isolates, but not in the control strain (Figure 6C). The mutation caused an early stop codon in two of three isolates (Clone 1, 3, Figure 6D). Transmission electron microscopy revealed complete absence of a capsule for the nonsense-mutated clones 1 and 3.

**Figure 6.**
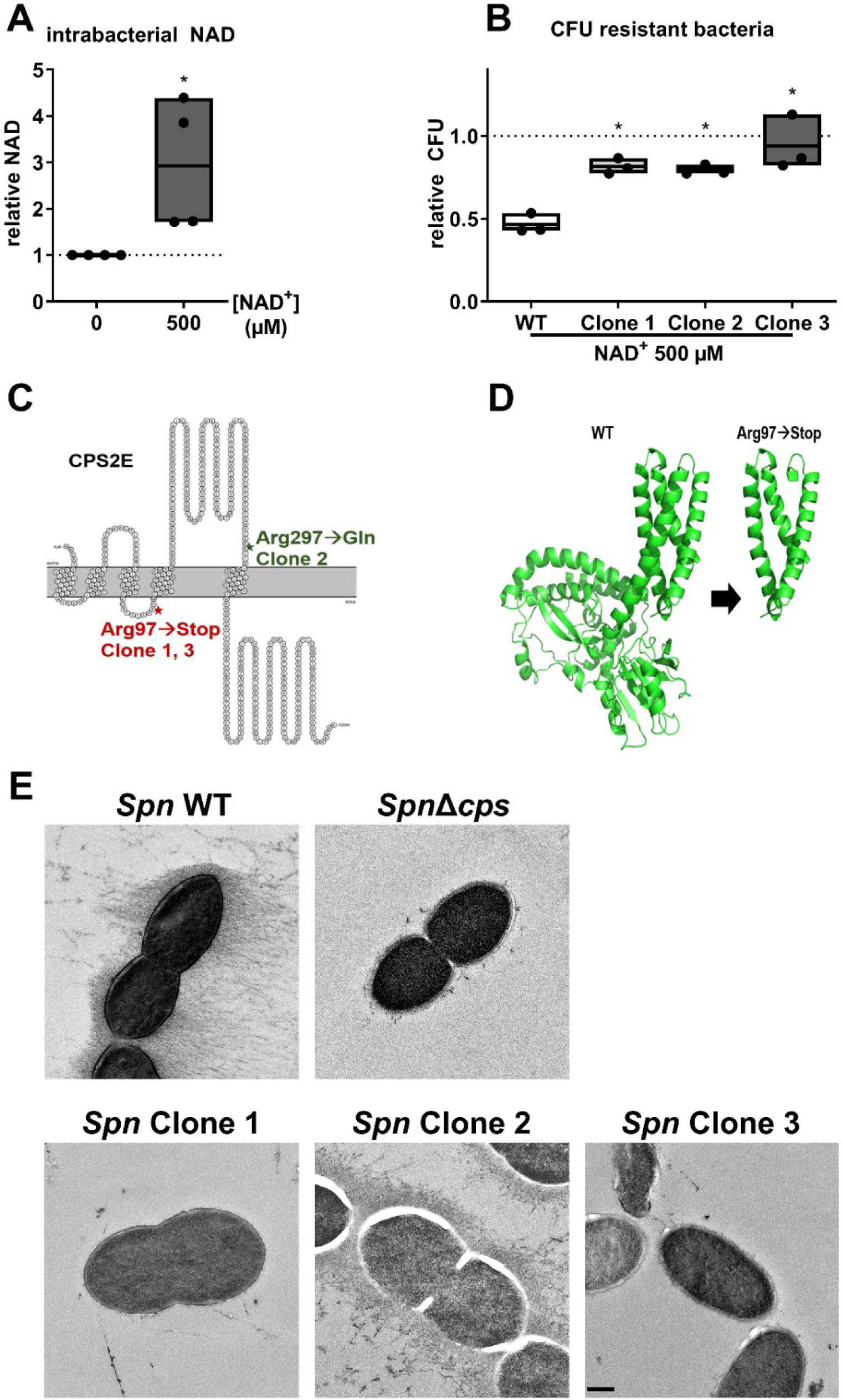
Development of NAD^+^ resistance is associated with loss of capsule. A) Spn was inoculated in cell culture medium with and without NAD^+^ treatment as indicated. Bacteria were lysed 6 h post-inoculation and the total intracellular NAD was determined. B-D) Spn was cultivated in cell culture medium and passaged 6 times with increasing concentrations of NAD^+^(50 µM to 5 mM) or without treatment. Afterwards, bacteria were plated on NAD^+^ (500 µM) supplemented agar plates. Individual clones were picked and characterized for NAD^+^ resistance and growth behavior and sequenced. B) Spn WT and resistant bacteria were treated with 500 µM NAD^+^ for 6 h and bacterial counts were determined. C) Spn clones passaged with increasing concentrations of NAD^+^ and a control passaged in cell culture medium without NAD^+^ were sequenced. Identified amino acid changes in the protein CPS2E are displayed. Asterisks indicate amino acid substitutions. D) 3D protein structure of CPS2E wild type and Arg977Stop, generated with AlphaFold. E) Transmission electron microscopy of Spn WT, Δcps and the NAD^+^ resistant clones. Bacteria were cultivated until early logarithmic phase in liquid media, cryo-fixated and the capsule stained using OsO_4_. (ns = not significant; * = p < 0.05; N = 3-4)

### 3.6. NAD^+^ acts antibacterial by interfering with the pneumococcal energy metabolism

To assess if capsule synthesis and NAD^+^ sensitivity are causally linked, we performed NAD^+^ treatment of a capsule deficient mutant of *Spn*. Treatment of *Spn*Δ*cps* with NAD^+^ for 9 h did not show any growth-limiting effects (Figure 7A). Biosynthesis of the pneumococcal capsule is an energetically costly process and closely linked to the bacterial energy metabolism (Echlin et al., 2016). Under energetically detrimental conditions, the capsule was shown to interfere with pneumococcal growth (Hathaway et al., 2012). We therefore hypothesized that NAD^+^ interference with intrabacterial ATP homeostasis is responsible for the growth limiting effect of NAD^+^. *Spn* WT, Δ*cps*, and NAD^+^-resistant clone 3 all showed a significant reduction in intrabacterial ATP after NAD^+^ treatment (Figure 7B). When left untreated, *Spn* clone 3 and *Spn*Δ*cps* showed an increase in intrabacterial ATP by a factor of 2 compared to the WT strain (Figure 7B). Treatment of bacteria with 5 mM of the ATP precursor pyruvate in addition to NAD^+^ treatment abolished the NAD^+^-dependent reduction in intrabacterial ATP and reduction in bacterial CFU (Figure 7C, D). In summary, the NAD^+^-mediated antibacterial effects were dependent on shifts in the bacterial energy metabolism and capsule biosynthesis.

**Figure 7.**
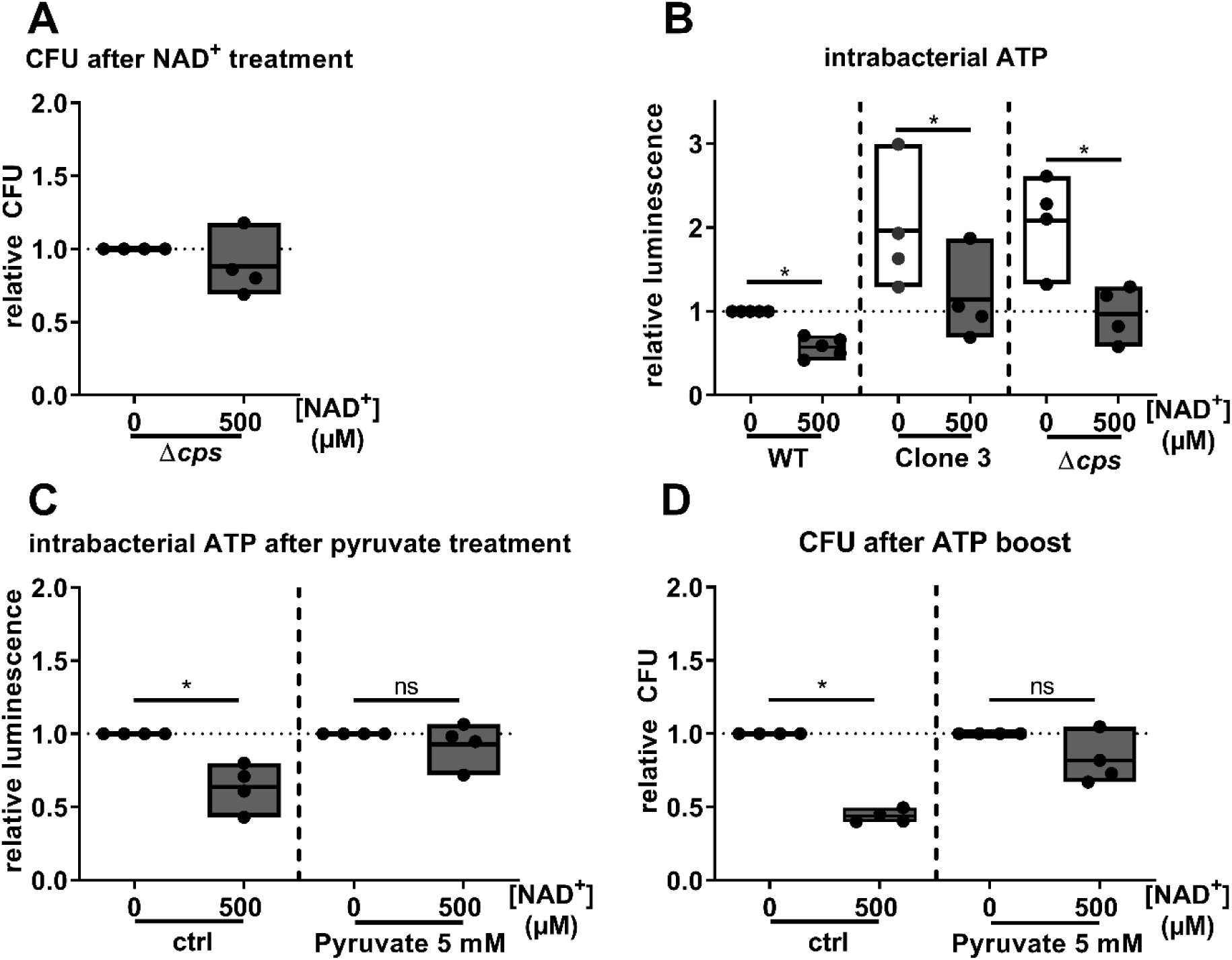
Development of NAD^+^ resistance is associated with metabolic adaptation. A) SpnΔcps was treated with NAD^+^ for 9 h or left untreated and bacterial counts were determined. Relative bacterial counts are depicted. B) Spn WT, clone 3 and SpnΔcps were incubated in cell culture medium for 6 h with and without NAD^+^ treatment. Afterwards, the intrabacterial ATP was determined by a commercial luminescence assay and normalized against bacterial counts. Measured luminescence is linear to intrabacterial ATP. C, D) Intrabacterial ATP after 6 h (D) and bacterial counts after 9 h (E) of combined treatment of Spn WT with 500 µM NAD^+^ and 5 mM pyruvate. Statistics: t-test (A, B); Two-way-ANOVA with Fisher’s LSD (C, D); Significance was determined against uninfected/untreated controls if not indicated otherwise. (ns = not significant; * = p < 0.05; N = 4)

## 4. Discussion

In this study, we describe a previously unreported interference of *Streptococcus*□*pneumoniae* with the host NAD metabolism. Our multi-omics approach revealed that in response to *Spn* infection, NAMPT is upregulated in bronchial epithelial BEAS-2B cells as part of a general, pro-inflammatory response. This is counteracted by a *Spn*-specific mechanism to downregulate NMNAT1 expression. The observed gene regulations seem to be pathophysiological relevant, as primary human bronchial epithelial cells cultivated at air–liquid interface, human lung explants and lungs of *Spn*-infected mice showed similar gene regulations. Since NMNAT1 catalyzes the final step of the salvage pathway, the identified gene regulations result in a reduced intracellular concentration of NAD^+^.

To our knowledge, we are the first to show *Spn*-mediated interference with the host NAD^+^ homeostasis. NAD^+^ was, however, shown to be a major player during other infections. During the viral infection process, NAMPT is routinely upregulated in a JAK-STAT dependent manner. This results in an increased NAD^+^ biosynthesis to fulfil the energetic demands of the antiviral enzymatic machinery (Wellen and Hotamisligil, 2005). To evade this defense mechanism, several viruses synthesize macrodomains which counteract the function of NAD^+^-consuming PARP by hydrolyzing ADP ribosylation sites of host proteins (Chen et al., 2011; Shang et al., 2021). Direct interference of viruses with NAMPT expression was suggested to be mediated by the induction of miRNA expression, which prevents NAMPT translation (Chen et al., 2013; Gong et al., 2022). During bacterial infections, several bacterial pathogens, among them *Shigella flexneri* and the lung pathogens NTHi and *Mycobacterium tuberculosis*, were shown to utilize host-produced NAD^+^ for their own energy metabolism (Mantis and Sansonetti, 1996; Kemmer et al., 2001; Boshoff et al., 2008). In accordance, in this study, NTHi infection induced NAMPT expression which might cause an increase in NAD^+^ production. Besides that, bacteria of the gut microbiota were shown to produce nicotinic acid as an NAD^+^ precursor to enhance host NAD^+^ production (Shats et al., 2020).

In contrast to these previously described upregulations of host NAD^+^ production by different bacteria, we here describe a *Spn* specific downregulation of the NAD^+^ biosynthesis via repression of NMNAT1 gene expression mediated by pneumolysin. To assess how this affects the infection process, we experimentally decreased NAD^+^ biosynthesis by knockdown of NAMPT or NMNAT1 prior to bacterial infections. Knockdown of NAMPT or NMNAT1 both reduced epithelial NAD production and increased bacterial replication. NAMPT is a multifaceted enzyme with previously reported cytokine function during pro-inflammatory processes (Galli et al., 2020). NMNAT1, in addition to its biosynthetic function, was reported to act as a chaperone (Zhai et al., 2006). To our knowledge, however, no involvement of NMNAT1 in infection processes has been reported before. To determine whether NAD^+^ biosynthesis or secondary effects of NAMPT and NMNAT1 are responsible for the observed effect, we treated *Spn* with NAD^+^ in various concentrations. This resulted in a host-cell independent reduction of bacterial replication, demonstrating a direct antibacterial effect of NAD^+^. To confirm that epithelial NMNAT1 can produce sufficient NAD^+^ to exert antibacterial activity, we supplemented host cells with the NMNAT1 substrate and NAD^+^ precursor NMN during infection. NMN was previously reported to increase intracellular NAD^+^ concentrations *in vitro* and *in vivo* (Yoshino et al., 2018; Igarashi et al., 2022). Measuring intracellular NAD^+^ following NMN treatment showed an increase by approximately a factor 2.5, underlining the feasibility of our approach. Strikingly, in the presence, but not in the absence of host cells, NMN reduced the replication of *Spn*, suggesting an involvement of host cell mediated biosynthetic processes in antibacterial effects of NMN. Next, we inhibited expression of NMNAT1 in BEAS-2B by siRNA transfection prior to infection and NMN treatment. Thereby, we suppressed the cells’ ability to utilize NMN for NAD^+^ synthesis. Following siNMNAT1 knockdown, intra-host cell concentration of NAD^+^ was reduced and bacterial replication partially rescued, confirming the role of NMNAT1 in antibacterial NAD^+^ production. Remaining growth-inhibitory effects of NMN might be due to remaining NMNAT1 or enzymatic activity of the related enzymes NMNAT2 and NMNAT3 (Emanuelli et al., 2001; Raffaelli et al., 2002). Additionally, NMN treatment of primary human bronchial epithelial cells cultivated at an air– liquid interface equally increased intracellular NAD and reduced bacterial replication, supporting relevance in physiological settings. Likewise, treatment of BEAS-2B cells with the NAMPT inhibitor FK-866 (Hasmann and Schemainda, 2003) and the NAMPT activator SBI-797812 (Gardell et al., 2019) resulted in increased and decreased *Spn* replication, respectively without requiring an extrinsic NMN boost. This demonstrates the intrinsic potential of the NAD^+^ salvage pathway to maintain antibacterial NAD^+^ concentrations.

While a previous study showed shifts in metabolic gene expression of *Spn* upon NADH treatment, no growth limiting effects were observed (Afzal et al., 2018). Besides that, the effect observed in this study is not limited to the *Spn* D39, as treatment of *Spn* TIGR4 with NAD^+^ equally inhibited bacterial growth. It is, however, not a general antibacterial mechanism, as *S.aga* and the host ATP-dependent NTHi (GILDER and GRANICK, 1947) showed improved growth upon NAD^+^ treatment. *Spn* frequently appears as an asymptomatic colonizer of the upper airways. The bacteria-specific responses to dysregulated NAD^+^ production raises the interesting question of whether and how NAD^+^ is involved in the interaction between *Spn*, airway microbiota and the host. In summary, boosting the host-cell NAD^+^ metabolism specifically inhibits pneumococcal replication during the infection process.

Since NAMPT was previously reported to be regulated via the JAK-STAT pathway (Dantoft et al., 2019; Huffaker et al., 2021), we hypothesized NMNAT1 to be regulated via a similar pathway. However, while in accordance to the literature inhibition of JAK signaling by Ruxolitinib (Mesa, 2010) reduced NAMPT expression after *Spn* infection, expression of NMNAT1 was not affected. In the following, we assessed which pneumococcal virulence factor is responsible for the observed gene regulation. Virtually all pathogenic strains of *Spn* express the pore-forming cholesterol dependent cytolysin pneumolysin (Canvin et al., 1995; Tweten, 2005). Besides its cytolytic effect, pneumolysin was shown to induce TLR4-dependent and-independent pro-inflammatory responses and cause DNA damage in the host cell (Zysk et al., 2001; McNeela et al., 2010; Rai et al., 2016). Deletion of pneumolysin from *Spn* D39 rescued NMNAT1 expression in BEAS-2B after infection, while NAMPT expression remained unaffected. Furthermore, treatment of BEAS-2B with purified pneumolysin in sublytic concentrations caused a dose-dependent downregulation of NMNAT1 and NAMPT. The effects were dependent on lytic activity, as a lytically inactive variant of pneumolysin did not affect NMNAT1 and NAMPT expression. Interestingly, *S.aga,* which produces another pore-forming toxin, caused a downregulation of NMNAT1 similar to *Spn* (Aroian and van der Goot, 2007). These results support the hypothesis of dysregulation of host NMNAT1 expression as a pneumococcal mechanism to inhibit detrimental NAD^+^ production in the host. In contrast, the observed downregulation of NAMPT during treatment with purified pneumolysin might be masked by the pro-inflammatory JAK-STAT dependent upregulation of NAMPT. Intracellular bacteria like *Mycobacterium tuberculosis* were previously reported to export toxins into the cytoplasm which deplete host-cell NAD^+^ and interfere with the host metabolism (Sun et al., 2015; Tak et al., 2019). Related bacteria like Group A *Streptococcus* were shown to express NAD^+^ glycohydrolases which deplete intra-host-cell NAD^+^ and might thereby defend themselves against NAD^+^ mediated bacterial killing (Hsieh et al., 2018). For a pore-forming cytolysin, to our knowledge, interference with the host cell energy metabolism has not been reported before.

To gain insight into the antibacterial mechanism of NAD^+^, we generated three NAD^+^ resistant strains of *Spn* and compared their genome sequence to the wild type. Resistance against NAD^+^ treatment was associated with resistance against NMN treatment in presence of host cells, further supporting NAD^+^ production as a direct antibacterial mechanism. Strikingly, all sequenced isolates showed mutations in the gene encoding for the protein CPS2E. CPS2E catalyzes an initial step in the attachment of the pneumococcal capsule to the membrane. Amino acid exchanges in this protein were previously shown to render *Spn* unencapsulated (James et al., 2013). The lack of capsule for two of the generated strains, both containing an early stop codon, was confirmed by electron microscopy. In line with this observation, an *Spn* strain with deleted *cps* locus was resistant against NAD^+^ treatment in our experimental setting. The pneumococcal polysaccharide capsule surrounds the whole bacterial cell and is a major virulence factor, protecting *Spn* against adverse environmental conditions and the immune defense (Paton and Trappetti, 2019). Capsule production, however, is a major biosynthetic effort and closely intertwined with the bacterial energy metabolism (Echlin et al., 2016; Echlin et al., 2020). Starvation of *Spn* leads to a reduction in capsule size (Hamaguchi et al., 2018) and under Cadmium-induced metabolic stress *Spn* was shown to reduce capsule production (Hamaguchi et al., 2018; Neville et al., 2020). We therefore hypothesized that abolishing capsule production is an adaptation to a dysregulated energy metabolism. To test our hypothesis, we measured intrabacterial ATP with and without NAD^+^ treatment. NAD^+^ treatment of *Spn* WT resulted in a reduction of intrabacterial ATP. Strikingly, an NAD^+^ resistant strain and *Spn*Δ*cps* showed an approximately two-fold increased amount of basal intrabacterial ATP, supporting the link between NAD^+^ resistance and ATP homeostasis. We then aimed to abolish the dysregulation of ATP production by treating bacteria with the ATP precursor pyruvate in parallel to NAD^+^ treatment. Indeed, pyruvate treatment increased intrabacterial ATP and abolished the antibacterial effects of NAD^+^. Collectively, these data highlight the link between NAD^+^ stress, capsule biosynthesis and bacterial metabolic dysregulation, and provide a mechanism for antibacterial activity of NAD^+^.

There are limitations to our study. It is difficult to determine how the NAD^+^ concentrations used in this study relate to the actual extracellular concentration of NAD^+^ *in vivo*. In human blood, NAD^+^ concentrations were determined to be approximately 33 µM (Yang et al., 2022). While this is below the threshold for a significant growth inhibition of *Spn* in our study, concentrations might be higher in the infection-relevant microenvironments like the lung alveoli, especially upon infection-induced host cell lysis (Adriouch et al., 2007).

Furthermore, extensive animal experiments to confirm increased NAD^+^ concentrations after NMN supplementation and its effect on bacterial replication were beyond the scope of this study. Nevertheless, we were able to confirm the NAD^+^ salvage dysregulation in infected mouse lungs and showed the antibacterial effects of NAD^+^ production in *ex vivo* human lung infection models. Since an increase of NAD^+^ levels following NMN treatment *in vivo* is well established (Nadeeshani et al., 2022), it seems likely that the antibacterial effect of increased NAD^+^ production by NMN supplementation will hold up *in vivo*.

Finally, this study investigated the effect of differentially regulated host NAD^+^ biosynthesis on the infecting bacteria. However, repressed NAD^+^ production is likely to have important consequences on host cell function as well. NAD^+^ is an important energy source for antiviral host responses (Wellen and Hotamisligil 2005) and is expected to be similarly important for mounting antibacterial responses. Furthermore, reduced intracellular NAD^+^ and therefore reduced activity of NAD^+^-dependent sirtuins was associated with aging processes (Zhu et al., 2022). It was recently proposed that residential bacteria contribute to Alzheimer disease and progression (Bulgart et al., 2020). Should the observed dysregulation of NAD^+^ production continue after bacterial clearance, this raises the fascinating question of whether pneumococcal infections, especially pneumonia and meningitis, cause long-term detrimental metabolic effects, contributing to accelerated lung or brain aging. The same question holds for other viral and bacterial infections that interfere with the expression of genes involved in NAD^+^ biosynthesis.

Taken together, our study reveals NAD^+^ biosynthesis as a previously overlooked antibacterial defense mechanism against *Streptococcus pneumoniae* and sheds further light on the importance of the energy metabolism as a key playing field of host–pathogen interaction during infection.

## 5. Funding Statement

This work has been funded in part by the Bundesministerium für Bildung und Forschung (Federal Ministry of Education and Research: PermedCOPD – FKZ 01EK2203A; ERACoSysMed2 – SysMed-COPD – FKZ 031L0140; e:Med CAPSYS – FKZ 01ZX1604E), the Deutsche Forschungsgemeinschaft (SFB/TR-84 TP C01), and the von-Behring-Röntgen-Stiftung (66-LV07) to B.S., as well as the Hessisches Ministerium für Wissenschaft und Kunst (LOEWE Diffusible Signals LOEWE-Schwerpunkt Diffusible Signals) to B.S, A.L.J., A.B. and E.M.E..

## 6. Acknowledgments

The authors wish to thank Manuela Gesell Salazar for conducting the mass spectrometric measurements and Kerstin Hoffmann and Peter Claus as well as the team from the Microscopy CORE Lab, Maastricht for their excellent technical assistance. Figure illustrations were created using biorender.com.

## 7. Author contributions

Conceptualization: BK, AW, EV, BB, BS; Methodology: BK, AW, KS, JS, KK, UV, EV, BB, BS; Formal analysis: BK, WB, JW, NP, JS, BB; Investigation: BK, AW, KS, SB, KK, IB, JS; Resources: AK, MA; Writing – Original Draft: BK, BB; Writing: Review and Editing. BK, AW, WB, NP, KS, JW, JS, EME, AK, KL, AJ, AB, UV, EV, BB, BS; Visualization: BK, EME, BB; Supervision: AB, UV, EV, BB, BS; Funding acquisition: BS

## 8. Ethics statement

Bronchial tissue for cell isolation was kindly provided by the Biobank platform of the German Center for Lung Research (DZL, Biobank Giessen, Hesse, Germany). Donated tissue was handled in accordance to local ethics regulations (Philipps University Marburg; permit number: AZ 224/12) and analysed anonymously.

## 9. STAR Methods

### Bacterial strains and culture

*Spn* strains D39*, D39*Δ*ply,* D39Δ*cps* and TIGR4 were kindly provided by Prof. Sven Hammerschmidt (University of Greifswald, Germany). A clinical isolate of *S.aga* obtained at the University Medical Center Marburg was kindly provided by Frank Sommer (Phillips University Marburg, Germany), whereas NTHi 19418 was obtained from ATCC. *Spn* and *S.aga* were plated on Columbia Blood agar plates (Becton Dickinson, 254005) and incubated over night at 37 °C / 5% CO_2_. Single colonies were inoculated into Todd-Hewitt-Yeast (THY) media and incubated at 37 °C / 5% CO_2_ until early logarithmic phase (OD_600_ 0.3-0.4). Bacteria were sedimented (3,000g, 15 min, RT) and adjusted to OD 0.1 in BEGM, corresponding to 5×10^7^ CFU per ml (*Spn*) or 2×10^7^ CFU/ml (*S.aga*), respectively. NTHi was plated on Chocolate Blood agar plates (Becton Dickinson, 257011) and incubated over night at 37 °C / 5% CO_2_. Single colonies were inoculated into Brain-Heart infusion (BHI) media, supplemented with Hemin and NAD^+^ and incubated at 37 °C / 5% CO_2_ until early logarithmic phase (OD_600_=0.3-0.4). Bacteria were sedimented (3000g, 15 min, RT) and adjusted to OD 0.1 in BEGM; corresponding to 5×10^7^ colony forming units (CFU)/ml. To generate NAD^+^ resistant strains of *Spn* D39, bacteria were cultivated in THY medium. After 3 h of incubation per step, they were continuously diluted to fresh medium with gradually increasing concentrations of NAD^+^ (Merck, N7004; 50 µM, 100 µM, 250 µM, 500 µM, 1 mM).

### Bacterial infection and cell stimulation

For BEAS-2B infection experiments, bacteria were inoculated into fresh BEGM (Lonza, CC-3170) medium according to desired multiplicity of infection (MOI). Alternatively, cells were treated with the TLR2 ligand lipoteichoic acid from *Staphylococcus aureus* (LTA, 1 µg/ml, Invivogen, tlrl-pslta), the pore-forming cytolysin pneumolysin (Ply), Ruxolitinib (10 µM, SelleckChem, S1378), FK-866 (1 µM, R&D Systems, 4808) or SBI-797812 (1µM, Merck, SML2791) whereas control cells were treated with the respective dissolvent.

### CFU assay and bacterial growth curve

For analysis of bacterial replication during infection, supernatant of infected cells was collected at indicated time points and 10-fold serial dilutions were streaked on blood agar plates. After incubation overnight at 37°C and 5% CO_2_, bacterial colonies were counted manually and CFU/mL were calculated.

To assess direct inhibitory effects of metabolites on *Spn*, 5×10^5^ CFU of *Spn* in early logarithmic growth phase were inoculated in BEGM and treated with NAD^+^ or NMN (Sigma-Aldrich, N3501). Bacterial cultures were cultivated at 37 °C / RT for indicated times and the CFU assay was performed as described above.

### Bacterial sequencing

Total DNA for sequencing was purified using a DNeasy Blood and Tissue kit (Qiagen). DNA libraries for sequencing were generated by applying a Nextera XT DNA Library Preparation kit (Illumina, FC-131-1024), and sequencing was performed on a MiSeq Desktop Sequencer (Illumina) using a MiSeq Reagent kit, version 3, for 2 75-bp paired-end reads (Illumina, MS-102-3001). At least 5.3 mio reads were obtained for each sample. The investigation for single nucleotide variants was carried out using the Basic Variant Detection tool (Qiagen, v.2.2) of CLC Genomic Workbench (Qiagen, v.21.0.3) with a minimum coverage of 10, minimum count of 8 and minimum frequency of 80 % for mapped reads.

### Electron microscopy

For electron microscopy, *Spn* strains were inoculated into THY medium and incubated at 37°C. When OD600 reached 0.35, the bacterial culture was centrifuged at 8,000 g for 10 min. A small pellet of cells was cryo-fixed between two golden 75 µm apertures in liquid ethane using the sandwich plunge freezing method (Baba, 2008) and freeze-substituted in 2% osmium tetroxide (Sigma, 201030), 0.1% uranyl acetate, and 5% distilled water in acetone using the fast low-temperature dehydration and fixation method. Cells were infiltrated overnight in Epon resin (LADD, NC9925769) and polymerized at 60°C for 48 h. 80-nm-thick sections were cut with a Leica UC6 ultramicrotome and imaged with a Tecnai T12 (FEI, Eindhoven) transmission electron microscope running at 120 kV.

### ATP measurement

Total concentration of intra-bacterial ATP was determined using a luminescence assay (Promega, G8230) according to manufacturer’s instructions. ATP concentrations were normalized to bacterial counts as determined by CFU assay.

### Purification of Pnneumolysin

LPS-free pneumolysin was purified from a recombinant Listeria innocua 6a strain expressing PLY. Lytically inactive pneumolysin was generated by deleting alanine at position 146 (Kirkham et al., 2006).

### Culture and transfection of BEAS-2B cells

The human bronchial epithelial cell line BEAS-2B (CRL-9609, ATCC) was routinely cultured in BEGM (Lonza, CC-3071) according to ATCC protocol without use of fibronectin coating. Cells were seeded at 10^5^ cells/cm^2^ and cultivated to 80% confluence (usually overnight). Medium was exchanged, followed immediately by infection or stimulation (see below). Transfection experiments were performed with 20 nM scramble control (4390843, Ambion), siNAMPT (Dharmacon, L-004581-00-0005) or siNMNAT1 (Dharmacon, L-008951-00-0005) in Lipofectamin RNAiMAX (diluted 1:100, Invitrogen). Lipofectamin and siRNA were mixed in OptiMem medium (ThermoFisher, 51985034) and incubated for 10 min at room temperature. From the transfection mix, 100 µl was used to transfect approximately 2×10^5^ cells in 400 µl BEGM (total volume 500 µl). Transfection was performed when seeding cells. After transfection, cells were cultivated for 48 h and used for infection experiments as described.

### Culture and infection of primary human bronchial epithelial cells

Primary human bronchial epithelial cells (HBEC) from healthy donors were obtained and cultivated in an air-liquid interface system as described previously (Karwelat et al., 2020). In brief, 6×10^4^ cells/cm² were seeded onto collagenized (Type 1 Collagen, Merck, CC050) Costar Transwell Permeable supports (3460, Corning) in Airway Epithelial Growth Medium (AEGM, Promocell, C-21160)). After reaching confluence, they were airlifted by aspiration of apical medium. Medium in the basolateral chamber was changed to differentiation medium (DMEM (ThermoFisher, 31966021)/AEGM (1:1) supplemented with 0.1 ng/mL retinoic acid (Merck) and penicillin/streptavidin). Cells were cultivated for 28 days and barrier formation was verified by measuring transepithelial electrical resistance (TEER). All donors developed a TEER > 880 Ω/cm². Cells were used at passages 3 or 5.

Basolateral medium was exchanged for differentiation medium without antibiotics on differentiation day 26, and again immediately before infection on day 29. To assess the effect of NMN on *Spn* growth, *Spn* were added apically at MOI 20 in 50 µl PBS. After one hour of incubation, cells were washed with 400 µl PBS. NMN 500 µM was added basolateral and cells were incubated for 16 h. For gene expression analysis, cells were apically infected with *Spn* at MOI 5 or 20 in 10 µl PBS without washing for 16 h.

### Preparation and infection of human lung tissue explants

Preparation and infection of human lung tissue explants was performed as described previously (Szymanski et al., 2012). Tissue was collected at the University Medical Centre Marburg by Andreas Kirschbaum in agreement with local ethics regulations (Marburg 161/17) after obtaining written informed consent from patients. In brief, tumor-distant, macroscopically tumor-free human lung tissue was obtained from tumor resections of bronchial carcinoma patients. Per tissue sample, multiple punch biopsies of equal size were generated and left uninfected or infected with *Spn* in quadruplicates for 12 h. Infections were done by injecting *Spn* into the tissue explants at 10^6^ CFU/160 mg lung tissue.

### Apoptosis measurement

BEAS-2B cells were detached by treatment with trypsin/EDTA for 4 min. Cells were stained with the Muse^®^ Annexin V and Dead Cell Assay Kit (Merck Millipore) according to manufacturer’s protocol, and measured on a Guava easyCyte 5HT flow cytometer (Merck Millipore).

### mRNA expression analysis

BEAS-2B cells were lysed in TRIzol™ (ThermoFisher, 15596026) and RNA was isolated by phenol-chloroform extraction. BEAS-2B Cells were lysed in trizol and RNA was isolated by phenol-chloroform precipitation. Explorative transcriptomic analysis was performed using DNA microarrays (Human G3 v2 Kit, 8×60k, Agilent technologies). Microarray data are published in GEO under the accession GSE195778 (https://www.ncbi.nlm.nih.gov/geo/query/acc.cgi?acc=GSE195778). In brief, purified total RNA was amplified using the Agilent Low Input QuickAmp kit, 200 ng labelled aRNA was hybridized following the Agilent protocol. Slides were scanned using the Innoscan 900 scanner (Innopsys, Carbononne France) at 2 µp/pixel and images were analyzed with Mapix 6.5.0. Raw data was processed in R using limma (Ritchie et al., 2015). Background correction was performed with the normexp model and spot intensities were quantile-normalized between arrays. Gene expression analysis by qPCR was performed as described previously (Karwelat et al., 2020). The following custom-made primers were used for qPCR: IL8: fwd: 5′-ACTGAGAGTGATTGAGAGTGGAC-3′, rev: 5′-AACCCTCTGCACCCAGTTTTC-3′; NAMPT: fwd: 5’-GGTTACAAGTTGCTGCCACC-3’, rev: 5’-AGCAAACCTCCACCAGAACC; NMNAT1: fwd: 5’-GTGATCTCCGGTAGCACTCG-3’, rev: 5’-CTTGGCCAGCTCAAACAACC-3’; NNMT: fwd: 5’-TAAGGAGATCGTCGTCACTG-3’, rev: 5’-CTGCTTGACCGCCTGTCTC-3’; RPS18: fwd: 5′-GCGGCGGAAAATAGCCTTTG-3′, rev: 5′-GATCACACGTTCCACCTCATC-3′. All samples were processed on a Quantstudio qPCR device (Life Technologies). Gene expression was calculated as ΔΔCT values and normalized towards mock-treated controls.

### Proteomic analysis

For SILAC (stable isotope labelling with amino acids in cell culture) standard generation, BEAS-2B cells were cultivated in DMEM with 2% FCS with heavy isotopes of lysine and arginine (EurisoTop) for at least three passages. For sample generation, cells were cultivated in DMEM with 2% FCS, without addition of labelled amino acids and control-treated or infected with *Spn* MOI 0.5 for 16 h as described above. Cells were then washed with PBS and harvested in solubilisation buffer (8 M urea, 2M thiourea in H_2_O). Protein extraction and processing was described elsewhere (Surmann et al., 2015). Each control or infected sample was mixed 1:1 with the marked standard and analysed by 1D gel electrophoresis (NuPAGE 4-12% Acrylamide Bis-Tris Medi Gel, Novex, Life Technologies) according to the manufacturer’s protocol. All bands were separately extracted from the gel and digested with 10 µg/ml trypsin, following peptide extraction and C_18_ purification (Merck Millipore). Five fractions of each sample were separated by nanoLC (Dionex UltiMate 3000, Dionex/ThermoFisher Scientific), ionized by TriVersa NanoMate (Advion, Ltd.) and measured by mass spectrometry (Q Exactive^™^ Hybrid-Quadrupole-Orbitrap, Thermo Fisher Scientific). By using Proteome Discoverer^™^ 1.4 (Thermo Fisher Scientific), detected peptides were mapped to the Uniprot protein database. Per sample and identified protein, a light-to-heavy (i.e., sample-to-standard) ratio was calculated. These ratios were used to further calculate log2 fold-changes (infected vs control) and q-values by moderated t-test (data analysis described previously in (Benedikter et al., 2019)). The mass spectrometry proteomics data have been deposited to the ProteomeXchange Consortium via the PRIDE partner repository (Perez-Riverol et al., 2022) with the dataset identifier PXD039059. Details on data acquisition are outlined in Table S1.

### Metabolite analysis

After 16 h of infection with *Spn* MOI 1, infected BEAS-2B cells or infected controls were washed with 0.9%, 37 °C NaCl solution. Cell culture plates were placed on ice and cells lysed in ice-cold Tris-EDTA/MeOH buffer (100 mM Tris, 1 mM EDTA in *A.dest*, 1:1 mixture with 98% MeOH). An equal volume of ice-cold chloroform was added and samples were incubated for 30 minutes at 4 °C prior to centrifugation (10,000 g,-10 °C, 10 min). Supernatants were filtered (Minisart RC4, 0.2 µM, Sartorius), snap-frozen in liquid nitrogen and stored at-80°C until analysis. Tris-EDTA/MeOH and chloroform were gassed with N_2_ immediately before use to prevent oxidation of metabolites. Quantitative determination of intracellular metabolites was performed using LC-MS/MS. The chromatographic separation was performed on an Agilent Infinity II 1290 HPLC system using a ZicpHILIC SeQuant column (150 × 2.1 mm, 5 µm particle size, 100 □ pore size) connected to a guard column of the same specificity (Merck) at a constant flow rate of 0.35 ml/min, with mobile phase A being 10 mM ammonium hydroxide in water adjusted to a pH of 9.8, and eluent B being acetonitrile (Honeywell) at 30°C. The injection volume was 2 µl. The mobile phase profile consisted of the following steps and linear gradients: 0–7 min from 90 to 55 % B; 7–10 min constant at 55 % B; 10–10.1 min from 55 to 90% B; 10.1–12.5 min constant at 90% B. An Agilent 6495 ion funnel mass spectrometer was used in positive mode with an electrospray ionization source and the following conditions: ESI spray voltage 1,500 V, nozzle voltage 500 V, sheath gas 400°C at 12 l/min, nebulizer pressure 30 psig and drying gas 250°C at 13 l/min. Compounds were identified based on their mass transition and retention time compared to standards. Chromatograms were integrated using MassHunter software (Agilent, Santa Clara, CA, USA). Absolute concentrations were calculated based on an external calibration curve prepared in sample matrix.

### ELISA and Western Blot

Concentration of IL-8 in the cell supernatant was analysed with a commercial ELISA kit (OptEIA, BD Biosciences) according to the manufacturer’s instructions. For Western Blotting, cells were harvested in RIPA buffer and cell debris removed by centrifugation (8000 g, RT). SDS-PAGE (10% polyacrylamide) and wet blotting to a nitrocellulose membrane was performed with 25 µg protein as determined by BCA assay. For NAMPT detection, rabbit anti-human NAMPT (dilution 1:500; Thermo Fisher Scientific, PA1-1045) and anti-rabbit-HRP (dilution 1:1,000, NEB, 5127 S) were used. Actin detection was performed using the Goat anti-human actin (dilution 1:1,000, SantaCruz, sc-1616) and anti-goat HRP (dilution 1:5,000, SantaCruz, sc-2020). Turnover of ECL substrate (GE Healthcare, 28980926) was detected on a chemo-luminescence imager (INTAS Science Imaging Instruments).

### NAD^+^ / NADH measurement

Total extracellular, intracellular and intrabacterial concentration of NAD (NAD^+^ and NADH) was determined using a commercial colorimetric assay (Abcam, ab65348) according to manufacturer’s instructions on a Tecan Infinite M200 PRO (ThermoFisher). Approximately 2×10^5^ BEAS-2B cells were infected with *Spn* for 16 hours as described above. After infection, cells were lysed and NAD^+^ / NADH concentrations were determined. Cell counts of infected and uninfected samples were determined for normalization. To calculate intracellular concentrations, a BEAS-2B volume of 2.2 pl/cell was assumed based on literature (Madanayake et al., 2013).

## 10. Supplemental Figures

**Figure S1.**
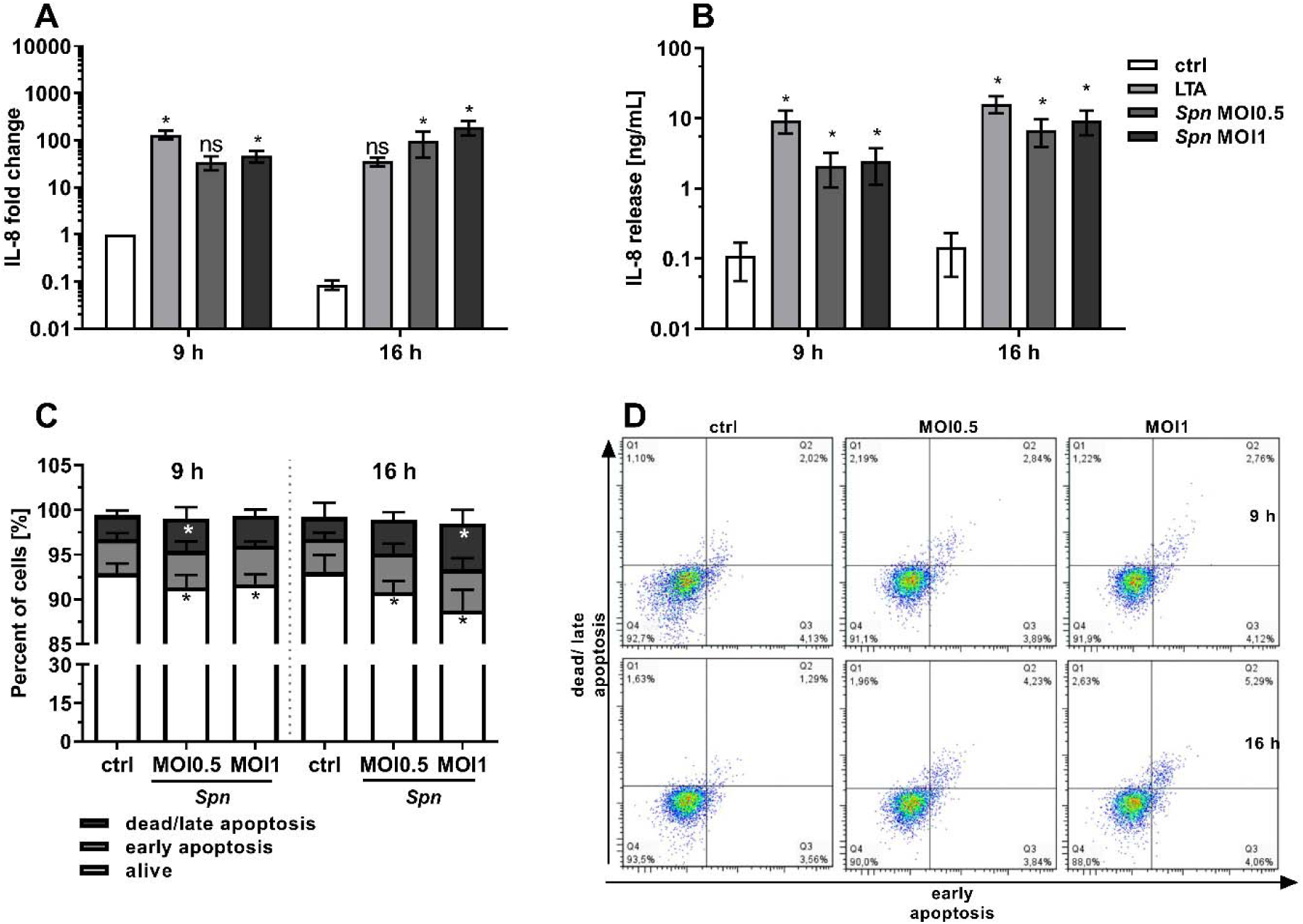
Spn infection of BEAS-2B induces a strong pro-inflammatory response with minimal apoptosis and cell death. BEAS-2B were left untreated, infected with Spn, MOI0.5 or MOI1 for 9 h or 16 h, respectively, or treated with LTA at 1 µg/ml. A) IL-8 gene expression as determined by qPCR. Results are normalized to the house keeping gene RPS18 and depicted relative to untreated control cells at 9 h. B) IL-8 protein secretion as determined by ELISA. C, D) Apoptotic state of infected BEAS-2B cells as determined by flow cytometry after AnnexinV/Propidium iodide staining. C) Quantification and D) scatter plot of stained cells. Statistics: Two-way ANOVA with Fisher‘s LSD. Significances were determined against uninfected controls if not indicated otherwise. (* = p < 0.05; N = 3).

**Figure S2.**
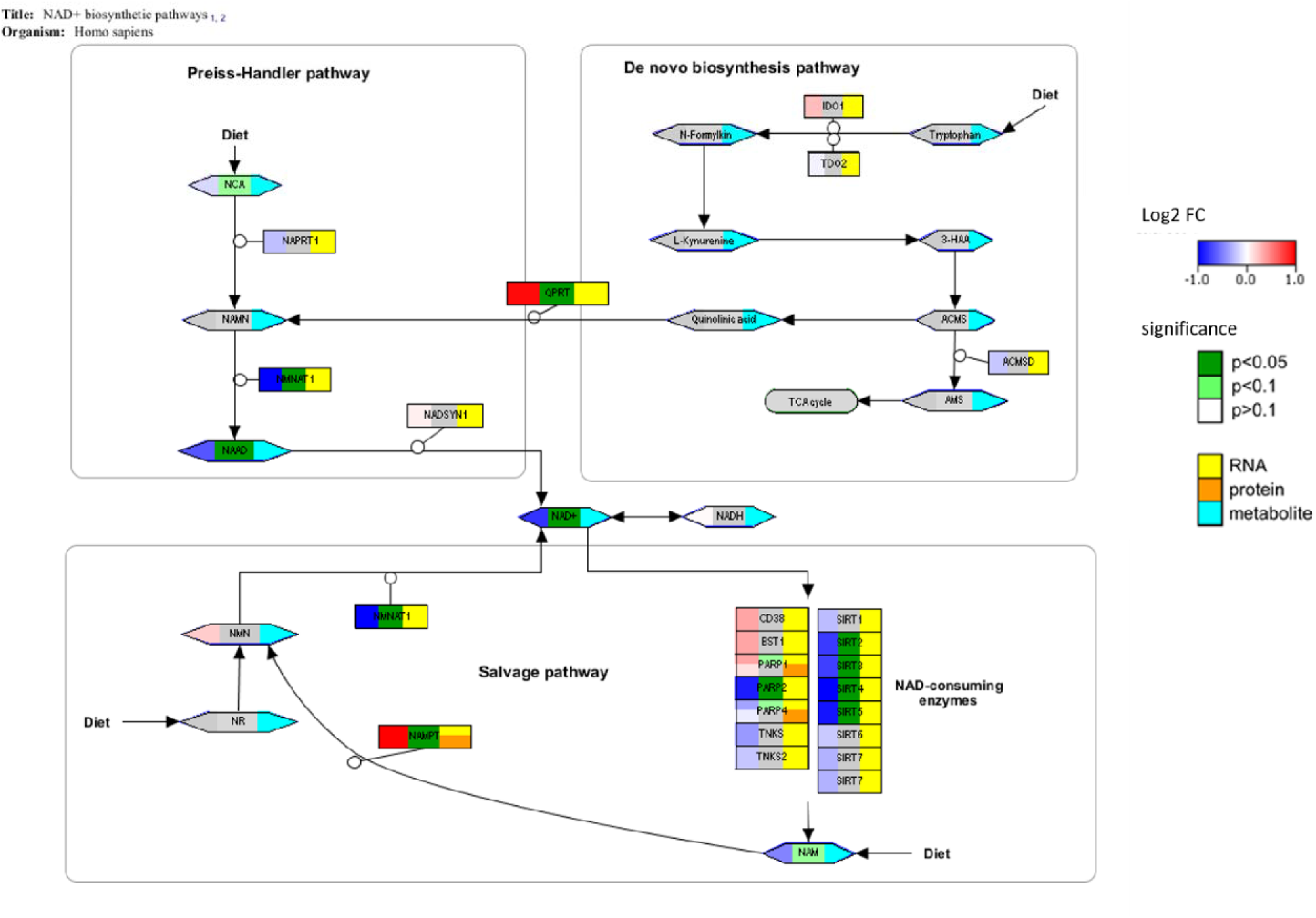
Regulation of NAD^+^ biosynthetic pathways upon pneumococcal infection. BEAS-2B lung epithelial cells were infected with Spn, MOI0.5 or MOI1 for 16 h. Host–cell proteome, transcriptome and intra-host–cell metabolite concentrations were measured and compared to uninfected controls. Multi-omics data was loaded into Pathvisio V3.3.0 (Kutmon et al., 2015) and visualized on the Wikipathways (Martens et al., 2021) Pathway WP3645 “NAD+ biosynthetic pathways” (https://www.wikipathways.org/pathways/WP3645.html).

**Figure S3.**
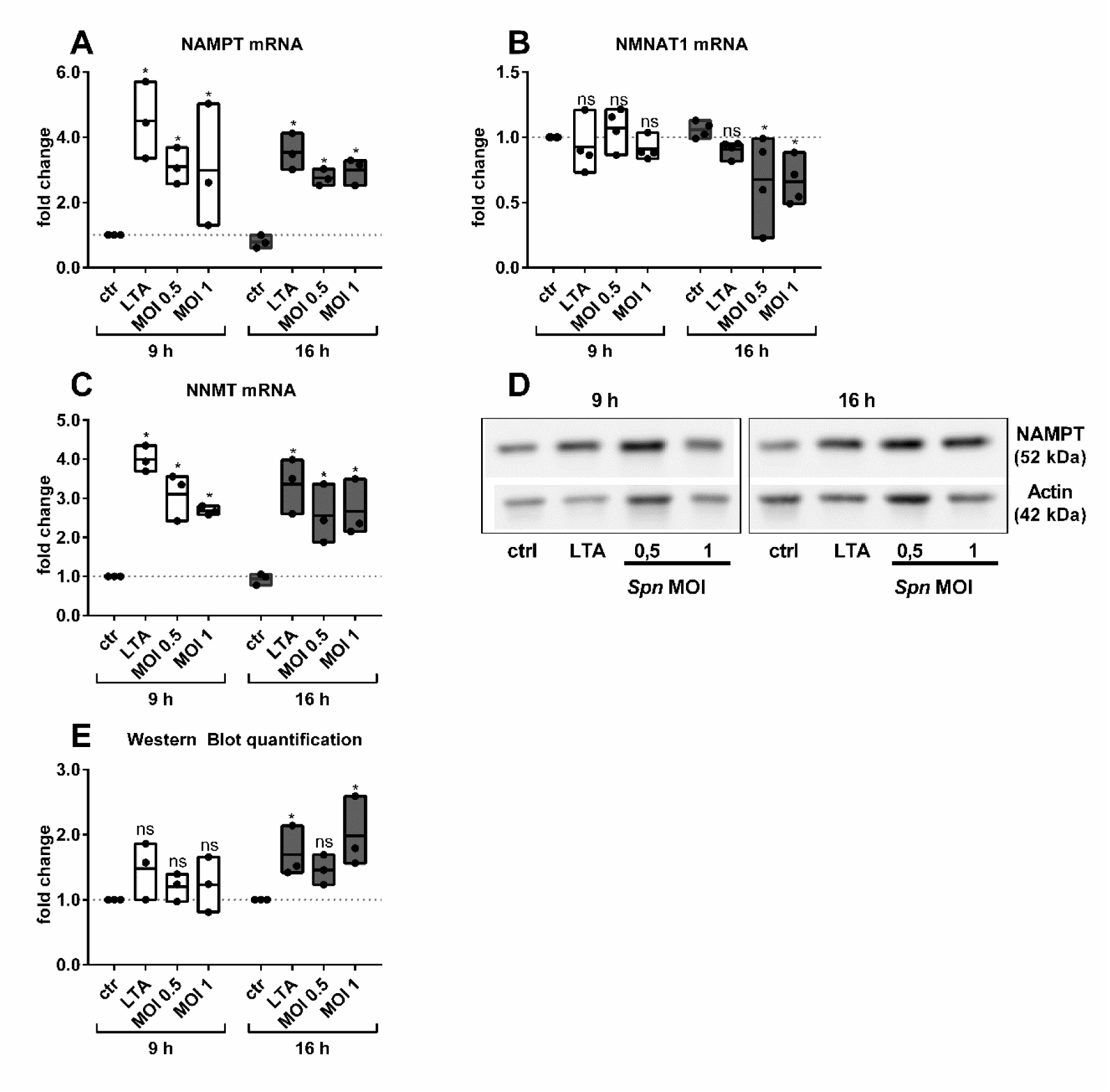
Regulation of NAMPT salvage-associated genes. BEAS-2B epithelial cells were infected with Spn, MOI 0.5 or MOI 1, treated with LTA (1 µg/ml) or left untreated for 9 h or 16 h, respectively. Afterwards, RNA and proteins were isolated. A-C) Expression of NAMPT (A), NMNAT1 (B) and NNMT (C) as assessed by qPCR Results are normalized to the house keeping gene RPS18 and depicted relative to untreated control cells.. D) Representative NAMPT Western Blot after stimulation/infection. E) Quantification of three biologically independent replicates of (D). Statistics: two-way ANOVA with Fisher‘s multiple comparisons. Significances were determined against uninfected controls if not indicated otherwise. (ns = not significant; * = p < 0.05; N = 3-4).

**Figure S4.**
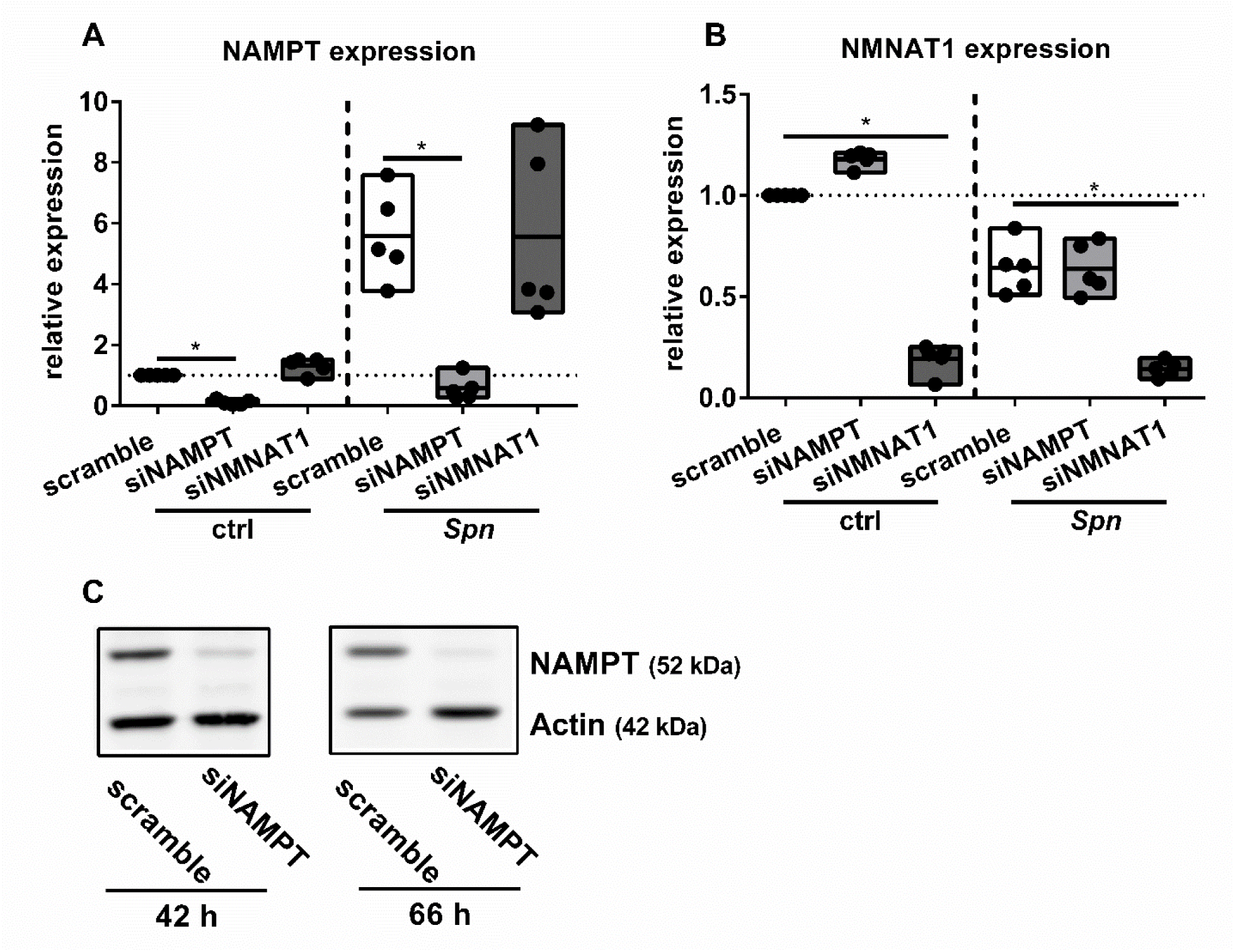
siRNA transfection induced a strong knockdown of NAD^+^ salvage genes. BEAS-2B were transfected with scramble, siNAMPT or siNMNAT1 as indicated. The medium was exchanged 48 h post transfection, and cells were either infected with Spn, MOI 1 for 16 h, or left untreated. A-B) RNA was isolated and qPCRs for indicated targets were performed. Results are normalized to the house keeping gene RPS18 and depicted relative to untreated control cells. C) Representative Western Blot analysis of NAMPT was performed after indicated time points. Statistics: t test; (* = p < 0.05; N = 5).

**Figure S5.**
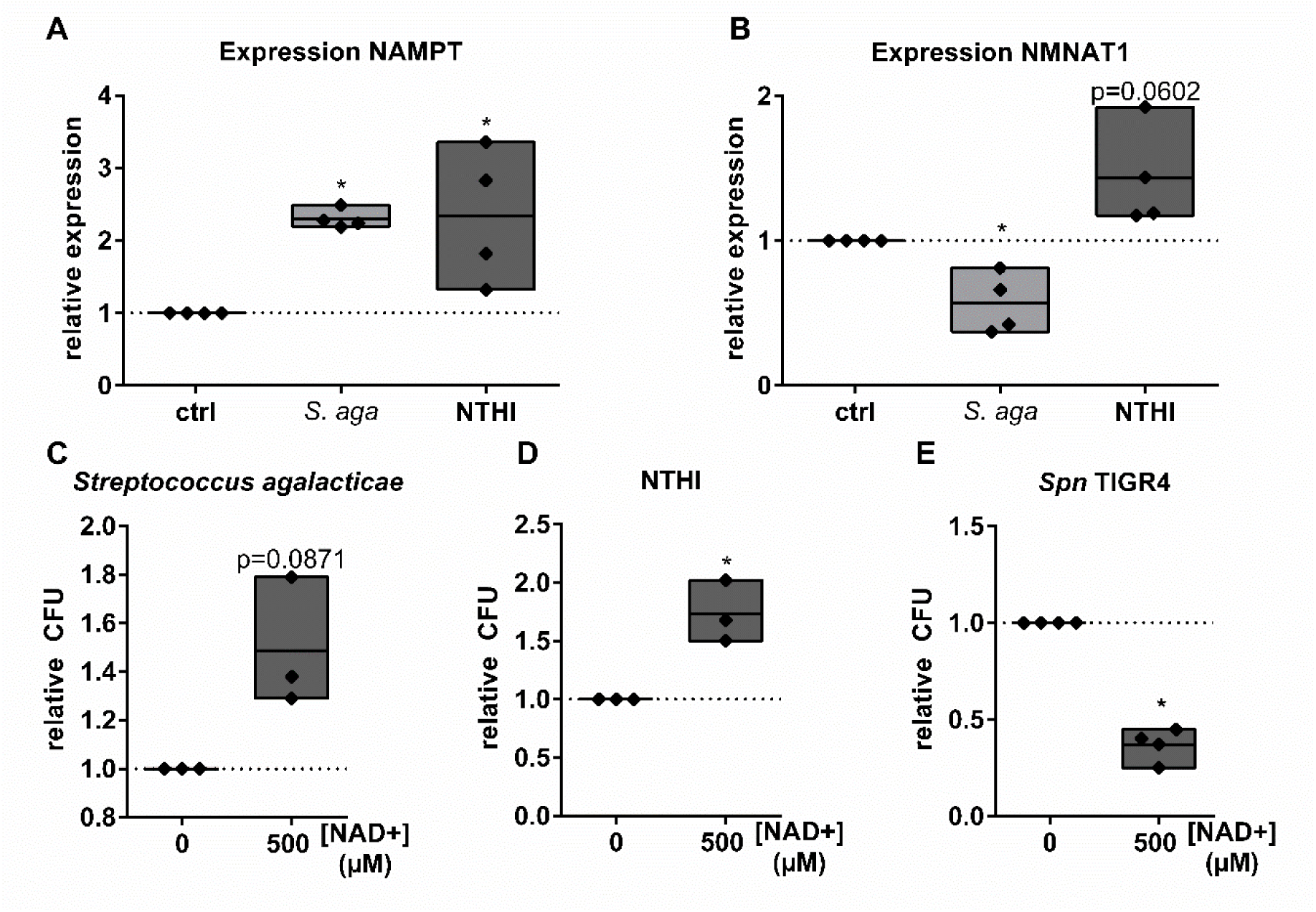
NAD^+^-mediated inhibition of bacterial replication and bacteria-induced regulation of NAD^+^ salvage genes are not general mechanisms. A-B) BEAS-2B were infected with NTHi or S.aga, MOI1, for 16 h or left untreated for control. Afterwards, RNA was isolated and expression of indicated NAD^+^ metabolic genes analyzed by qPCR. C-E) S.aga, NTHi and Spn TIGR4 were cultivated in BEGM for 6 h (TIGR4, S.aga) or 4 h (NTHi) and treated with 500 µM NAD^+^. Afterwards, bacterial replication was assessed by CFU assay. Statistics: t-test. (* = p < 0.05; ns = not significant; N = 3-4).

**Figure S6.**
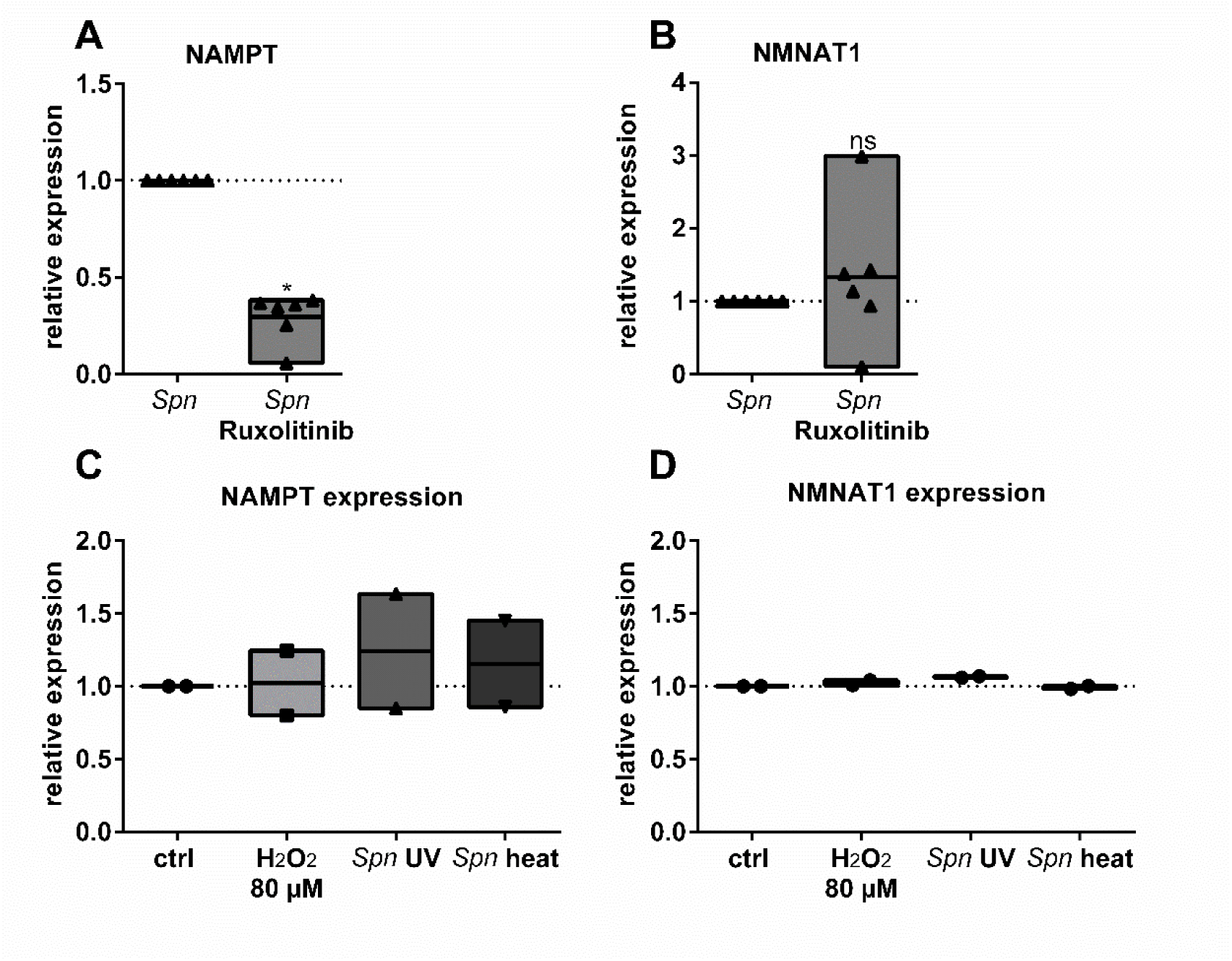
NAMPT and NMNAT1 are regulated by distinct mechanisms. A, B) BEAS-2B were infected with Spn with or without Ruxolitinib (10 µM) treatment for 16 h. RNA was isolated and expression of NAMPT and NMNAT1 determined. C, D) BEAS-2B were treated with 80 µM H_2_O_2_, UV-inactivated or heat-killed bacteria for 16 h. RNA was isolated and the expression of NAMPT and NMNAT1 determined. Results are normalized to the house keeping gene RPS18 and depicted relative to untreated control cells. Statistics (A+B): t-test; Significances were determined against untreated controls if not indicated otherwise. (* = p < 0.05; N = 4).

## Notes

### Competing Interest Statement

The authors have declared no competing interest.

